# SureMap: Versatile, Error Tolerant, and High Sensitive Read Mapper

**DOI:** 10.1101/173740

**Authors:** MohammadJavad Rezaei Seraji, Seyed Abolfazl Motahari

## Abstract

SureMap is a versatile, error tolerant and high sensitive read mapper which is able to map “difficult” reads, those requiring many edit operations to be mapped to the reference genome, with acceptable time complexity. Mapping real datasets reveal that many variants unidentifiable by other mappers can be called using Suremap. Moreover SureMap has a very good running time and accuracy in aligning very long and noisy reads like PacBio and Nanopore against reference genome.

## Background

After quality control of reads, workflows for processing of Next Generation Sequencing (NGS) datasets usually starts with mapping of reads to a reference genome [1]. In DNA resequencing, as an example, reads are mapped to a reference genome and variants are called to detect the differences between reference and target genomes. Undoubtedly, the quality of the read mapping step impacts our confidence of the final results of any downstream analysis. In particular, applying the results in diagnostics or therapeutics of any disease requires very reliable/provable steps in the workflows.

Read aligners handling different type of reads are in abundance. It may seem that there is no need for another aligner. This paper reveals the contrary: there are many situations where leftover reads that cannot be handled by current aligners contain very valuable information important in biology and medicine. These reads require special attentions and need more elaborate aligners to be handled. Moreover, the current technology is toward producing very long and noisy reads. Efficient handling of such reads is also a challenge requiring novel methodologies in read mapping.

Ideal read mapping amounts to finding the true origins of all reads. Three important factors inhibit achieving such a daunting task. First, the reference genome is usually very repetitive [2]. Over fifty percent of the human genome consists of repetitive elements such as L1, L2, and Alu elements [3, 4]. Second, reads are noisy. A perfect sequencing machine that outputs for each nucleotide its matching symbol does not exist. All machines create erroneous bases including mismatches, insertions, and deletions. Third, there exist variations between reference and target genomes. A read from a given location on a target genome may look more closer to somewhere else on a reference genome.

There exists many read mapping packages suitable for short/long/very long reads such as Bowtie [5], Bowtie2 [6], BWA [7, 8], SOAP [9], SOAP2 [10], MAQ [11], mrsFAST [12], Stampy [13], and Meta-aligner [14]. A single location or a list of locations of possible origins of each read is reported by these packages. In the popular *unique* map strategy, all reads mappable to more than one location are removed from the set of reads. This may leave many low coverage areas that causes, in turn, incorrect decision making.

Alternatively, a *perfect* read mapper can output all possible mapping locations and pass the information the the given downstream package that can handle such information, c.f., [15]. Theoretically, a perfect read mapper has no loss of information if it reports for each read a list of candidate locations where the true location always belongs to it. The shorter the list size, the closer we are to the ideal read mapper.

In this paper, we present *SureMap*, a fast, error tolerant, and very high sensitive read mapper. The mapper is able to map short and long reads to a reference genome and outputs a list of all possible locations within any given edit distance on the reference genome. SureMap is also able to map “difficult” reads with acceptable time complexity: those requiring many edit operations to be mapped to the reference genome. SureMap can be applied in many instances as follows.

First, some of the reads that cannot be mapped to a reference genome by current aligners contain very useful information that needs to be extracted. For instance, if the variations between a reference and target genomes are considerably high within a certain region then a more sensitive aligner with provable accuracy is needed to map the reads to the region and extract information regarding variations between the two genomes.

Second, it is very useful to evaluate the accuracy of calling variations by yet another strategy. In this way, the confidence in applying results in any downstream analysis increases significantly. For instance, using a conventional workflow one can obtain several candidate locations for variants. SureMap can be used to remap reads involved in calling these variants to measure the confidence of the calls. Presumed unique reads used to call a variant by a conventional aligner are not unique if they are mapped by SureMap.

Third, long-read sequencing machines such as PacBio RS and MinION produce reads that are very noisy with high indel errors. Mapping these reads to a reference genome is very challenging. SureMap can be used to align fragments of these reads to the reference genome such that the correct position of the reads can be inferred from the mapped fragments. SureMap is also tailored to align such very noisy reads incorporating ideas from Meta-aligner [14].

## Results

In this section, we evaluate the performance of SureMap with other read aligners with respect to speed and accuracy. Used in its highest sensitive mode, SureMap is usually slower than other less sensitive aligners. However, the cost paid for sensitive alignment is compensated with better and more accurate downstream analyses. On the other hand, SureMap, used in its heuristic mode, is fast and accurate in aligning very long reads. Using simulated and real datasets, the experimental results supporting the claimed performance are presented as follows.

### Simulated Datasets

We have compared SureMap with Bowtie [5] and SOAP2 [10] on aligning short reads. To this end, we have simulated 10M reads of length 50 from human genome hg19 with different error rates. The type of errors in the reads is mismatch errors. The simulated reads are mapped to the reference genome using 20 parallel threads. Due to the indexing schemes used in all three packages, the memory usages for Bowtie, SOAP2, and SureMap are, respectively, 2.4GB, 5.4 GB, and 8.7 GB.

#### Unique Alignment

Many down stream analysis throw away any read mapped to multiple locations on the reference genome. Therefore, a mapper that can report higher unique alignment within some given edit distance is preferable in downstream analysis. This is due the fact that removing non-unique reads will result in low coverage of some regions which in turn lowers the confidence of the statistical inference.

Figure 1 shows the result of mapping reads to the human genome using Bowtie and SureMap. Three different mismatch read error rates are used for the sake of comparison. For the error rate of 1%, Bowtie and SureMap are compared each allowing at most three mismatches. Both perform equally in mapping reads to the reference genome. For the error rate of 5%, SureMap outperform Bowtie in aligning unique reads by allowing 5 mismatches. For the error rate of 10%, SureMap can still uniquely map 70% of the reads to the reference genome by allowing 7 mismatches. This is a very interesting result as it shows that a low cost, high error rate, short read sequencer still produces enough information for mapping 70 percent of the reads.

**Figure 1.**
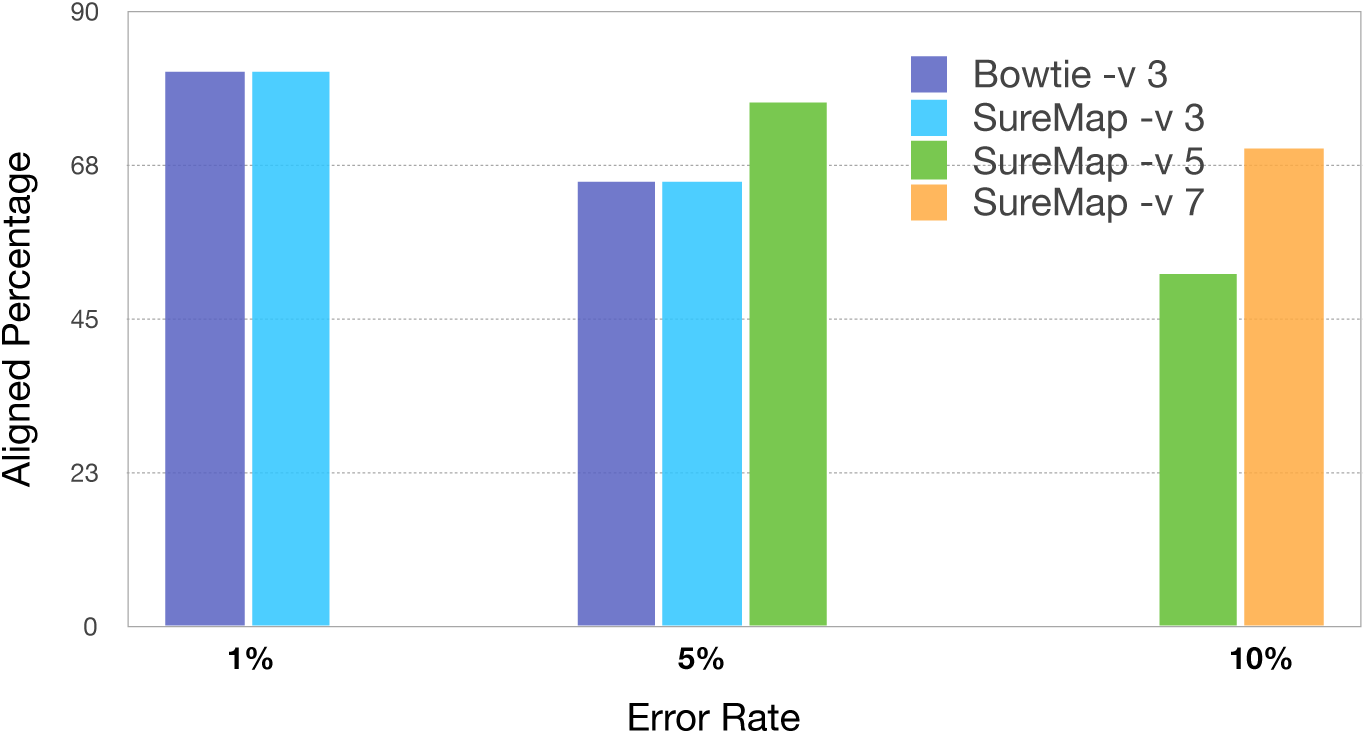
SureMap and Bowtie are compared with respect to percentage of uniquely aligned reads. The performance of the two aligners for read error rate 1% is the same. For higher read error rates SureMap is capable of uniquely align more than half of the reads if its highest sensitive option is chosen.

#### List Decoding

In Table 1, SureMap is compared with Bowtie and SOAP2 with respect to sensitivity and speed. For each read, up to 10 locations that meet the required threshold for the number of mismatches are reported. The table shows that the higher sensitivity achieved by SureMap is obtained by losing the speed. This reveals that SureMap is preferable to be used for reads that other aligners cannot handle them due to their high error/variation rates.

**Table 1.** Comparison between three aligners with respect to sensitivity and speed. 10 Mega Reads with length 50 are simulated from the human genome hg19 with different error rates.

| Error Rate | Aligner | Time (s) | Read aligned |
| --- | --- | --- | --- |
| 1 % | Bowtie -v 3 | 500 | 99.83% |
|  | SOAP2 -v 3 | 500 | 98.65 % |
|  | SureMap -v 3 | 700 | 99.83 % |
| 5 % | Bowtie -v 3 | 480 | 76.9% |
|  | SOAP2 -v 3 | 210 | 55.65 % |
|  | SureMap -v 3 | 600 | 76.9 % |
|  | SureMap -v 5 | 7500 | 96.3 % |
| 10 % | SureMap -v 5 | 10100 | 63 % |
|  | SureMap -v 7 | 16500 | 88 % |

### Real Datasets

#### Variant calling

In population scale projects such as the 1000-genome project [16], common and rare variants are identified to be used in association studies. In its basic pipeline, reads from a target genome is mapped uniquely to a reference genome and variants are called based on the mapped reads to a given genomic location.

There exit situations where sequencing reads are almost perfect but variations between two genomes are clustered within several genomic locations. Variant calling in these regions is very hard as reads cannot be mapped accurately to them. SureMap is specially useful to map reads to these regions and identify these nearby variants.

In this experiment, a sample from the 1000-genome project with the accession number SRR099983 is selected. Paired reads from this sample are mapped to the reference genome using BWA MEM and Bowtie2 in its very sensitive mode. It is possible to map 95% of the reads to the reference genome. The remaining 5% of reads (about 12 million reads) are mapped by SueMap using the unique option. SureMap could map only 2.5% of these reads amounting to 300,000 reads. Surprisingly, some of the mapped reads are clustered across the genome enabling a variant caller to identify new variations between reference and target genomes. In particular, using VCF tools with depth at least 8 and quality above 80 one can identify 500 new variations.

In Figure 2, the results of mapped reads are visualized using Integrative Genome Viewer [17, 18]. It is evident that calling the variants is impossible with either Bowtie2 or BWA MEM as the coverage is very low. SureMap, on the other hand, is able to identify all the variants with high confidence. Other mappers such as [19, 20, 21] could not handle these “difficult” reads as well.

**Figure 2.**
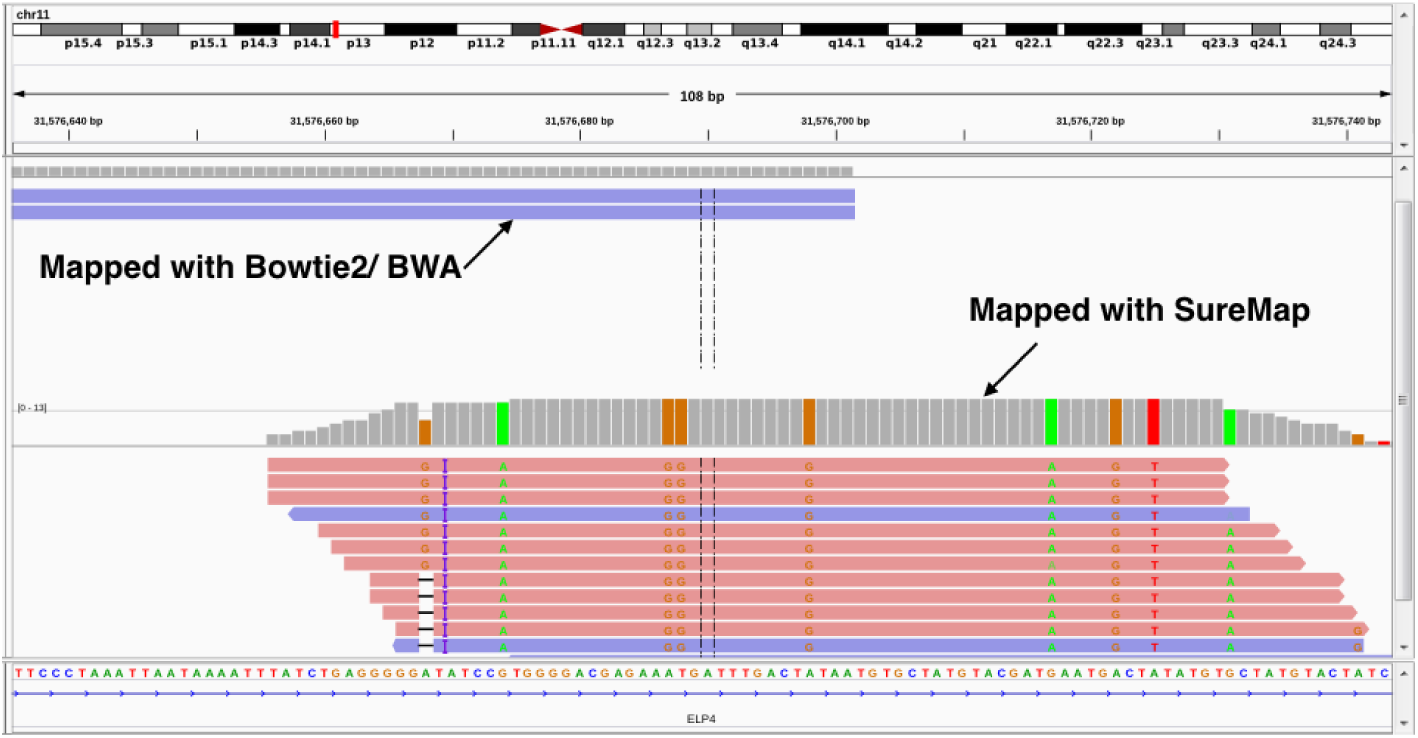
Reads from the 1000 genome project are mapped to the human reference genome using Bowtie2 (very sensitive mode) and BWA MEM. The remaining reads are mapped with SureMap where several new variants are identified. In this figure, a genomic location on Chromosome 11 is shown where reads mapped with SureMap are clustered together identifying 10 new variants.

#### Long Reads

To evaluate the performance of SureMap on very long reads, real datasets from two different platforms are selected: Pacbio and Nanopore.

From Pacbio, 25249 reads with total length 180 Mbp are extracted from the dataset http://datasets.pacb.com/2013/Human10x/READS/index.html (M130929, subreads 0). Reads are mapped to the human reference genome hg19 using a 32-core machine (30 threads are used for alignment) using four different aligners: SureMap, Bwa MEM, Blasr [22], and LAST [23]. The results are presented in Table 2. Depending on the length of fragments used in the local alignment in SureMap, six different modes of operations are considered for it, namely *ℓ* ∈ {200, 300, 650,1100,1600, 3000}. Bwa Mem is used in its “pacbio” modes and the other two aligners are used in their default modes. The second column of the table represents the running time of the aligners. To report the fraction of reads that are aligned to the reference genome, we have used four different filters depending on the rate of variations between reads and the aligned locations. Comparing SureMap with Bwa MEM, we observe that when *ℓ* = 650 SureMap is faster with higher quality in read mapping with little loss in coverage. SureMap outperforms Blasr with respect to time with little loss in quality and coverage. It is worth mentioning, Blasr in 5 percent of cases results in overlapping aligned fragments which is not desirable in downstream analyses. SureMap also outperforms LAST in time and total coverage and also in mapping the reads with low error rate. the memory usages for SureMap, bwa-Mem, blasr and LAST are, respectively, 8.9GB, 5.4 GB, 14.5 GB and 12.5 GB.

**Table 2.** The results of aligning PacBio reads with four different aligners: SureMap, Bwa MEM, Blasr, and LAST. Dataset source: datasets.pacb.com/2013/Human10x/READS/index.html (M130929, subreads 0). A set of 25249 reads with total length180 Mbp is sampled from the genome and mapped to the reference genome hg19. *ℓ* stands for the length of fragments used in SureMap. *v* denotes the variation rate between read and the reference genome. For example, Column *v* ≤ 0.3 shows the percentage of total reads (25249 reads) that an aligner program can align so that the variation rate between the aligned reads and their aligned region in the reference genome is less than 0.3. The last column which is called coverage represents the total length of all aligned reads with variation rate *v* ≤ 0.3.

|  | PacBio Table |  |  |  |  |  |  |
| --- | --- | --- | --- | --- | --- | --- | --- |
| | Time | $v \leq 0.3$ | $v \leq 0.25$ | $v \leq 0.2$ | $v \leq 0.15$ | Running Option | Coverage (Mbp) |
| SureMap | 95 s | 85.5 | 79.8 | 68.2 | 44.9 | $\ell=200$ | 130.0 |
| | 107 s | 88 | 82.2 | 70.5 | 46.3 | $\ell=300$ | 138.8 |
| | 162 s | 90.8 | 84.9 | 72.3 | 47.2 | $\ell=650$ | 150.7 |
| | 251 s | 91.37 | 85.2 | 72.5 | 45.1 | $\ell=1100$ | 154.1 |
| | 365 s | 92.0 | 85.6 | 72.5 | 47.2 | $\ell=1600$ | 156.5 |
| | 760 s | 92.4 | 85.4 | 72.5 | 48.0 | $\ell=3000$ | 159.1 |
| Bwa MEM | 180 s | 90.9 | 84.0 | 70.7 | 46.1 | -x pacbio | 159.8 |
| Blasr | 755 s | 94.6 | 90.1 | 77.8 | 51.6 | default | 161.6 |
| LAST | 751 s | 94.4 | 90.4 | 72.1 | 38.3 | default | 148.5 |

From Nanopore, 30143 reads covering 176Mb of the genome are extracted from the dataset ERR1676721. The same simulation setup is considered as that of PacBio reads except that Bwa MEM is used in its “ont2d” option and LAST is removed from the set of aligners since its time complexity was not in the order of other cases. The results are presented in Table 3. SureMap is able to map faster than the other two aligner with little loss in the number of mapped reads and total coverage.

**Table 3.** The results of aligning PacBio reads with three different aligners: SureMap, Bwa MEM, and Blasr. Dataset name: ERR1676721. A set of 30143 reads with total length 176 Mbp is sampled from the genome and mapped to the reference genome hg19. *ℓ* stands for the length of fragments used in SureMap. *v* denotes the variation rate between read and the reference genome.

|  | Nanopore Table |  |  |  |  |  |  |
| --- | --- | --- | --- | --- | --- | --- | --- |
| | Time | $v \leq 0.3$ | $v \leq 0.25$ | $v \leq 0.2$ | $v \leq 0.15$ | Running Option | Coverage (Mbp) |
| SureMap | 187 s | 90.0 | 87.1 | 80.0 | 61.2 | $\ell=200$ | 145.4 |
| | 204 s | 91.9 | 89.5 | 82.8 | 63.3 | $\ell=300$ | 155.8 |
| | 230 s | 93.1 | 91.1 | 84.8 | 65.0 | $\ell=650$ | 164.1 |
| | 307 s | 93.3 | 91.3 | 85.2 | 65.3 | $\ell=1100$ | 165.0 |
| | 411 s | 93.4 | 91.3 | 85.2 | 65.3 | $\ell=1600$ | 165.4 |
| | 700 s | 93.6 | 91.4 | 85.3 | 65.4 | $\ell=3000$ | 166.1 |
| Bwa MEM | 441 s | 94.4 | 91.7 | 84.3 | 64.1 | -x ont2d | 168.0 |
| Blasr | 615 s | 94.2 | 92.9 | 88.1 | 67.2 | default | 172.0 |

It is worth mentioning that in most cases SureMap reports the exact alignment of the whole read to the reference genome by creating the CIGAR symbol for each read. This is in contrast to many aligners where fragments of the reads are clipped for easy and fast alignment of the reads.

## Conclusion

SureMap is designed and implemented to handle reads requiring special attention during read mapping. SureMap is very sensitive and versatile: it can handle short, long, and very long reads alike. Embedded in SureMap is a perfect list reporter that for a given read it reports all possible locations on the reference genome that are within a predefined edit distance to the read. SureMap is used in practice to find many interesting results unraveling its capability in handling special reads. Reads from the 1000 genome project not mappable with other aligners can be mapped to the reference genome with SureMap and new variants can be identified. SureMap uses a very flexible algorithm that works well for very long and noisy reads and at the same time it can make a perfect balance between speed and accuracy. For example, running fast mode of SureMap for Pacbio datasets, we can see that it is almost 2X faster than bwa-MEM while its coverage is no less than 80 percent of bwa-MEM mapping coverage.

## Methods

Mapping a read *R* to a given reference genome *X* is an instance of the well known approximate string matching problem [24, 25]. In fact, we aim at finding all substrings of the genome within edit distance *d* from *R*. More formally, the list of substrings is represented by

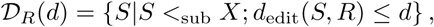

where *S* <_sub_ *X* means that *S* is a substring of *X*. When the read and the edit distance are clear from the context, 𝓓_*R*_(*d*) is simply replaced by 𝓓.

### Divide and merge

The concatenation of two strings *A* and *B* is denoted by *c*(*A*, *B*). For a given *S* ∈ 𝓓_*R*_(*d*), there exists a partition of *S* = *c*(*S*^L^, *S*^R^) and a bipartition of *R* = *c*(*R*_1_, *R*_2_) such that

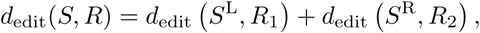

where *R*_1_ and *R*_2_ are, respectively, the first and second halves of the read *R*. This simply implies that either 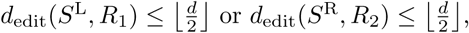 for a given threshold *d*. Next, we can define two new sets:

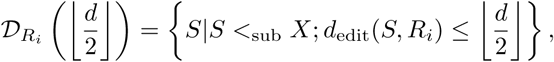

where *i* ∈ {1, 2}. In this way, we arrive at two new problems where in each of them the read length as well as the edit distance are halved. These are supposedly simpler sets to be listed by a mapper.

The basic strategy in our algorithm is based on the preceding partitioning of the reads. Let us assume that we have found the set 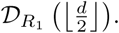 For each *S*^L^ ∈ 𝓓_*R*_1__, we start extending *S*^L^ base by base to the right on the genome and fill out a dynamic programming table to obtain the minimum edit distance between *R* and a substring of the genome starting at *S*^L^. If we can find *S* = *c*(*S*^L^, *S*^R^) within *d* distance from *R*, then we include *S* in 𝓓_*R*_(*d*). A similar procedure can be performed on 𝓓_*R*_2__. This time, however, we need to extend the substrings in 𝓓_*R*_2__ to the left.

In this way, we obtain the set 𝓓. However, some substrings obtained from the left extension can also be obtained by the right extension. To avoid this, we note that once we obtain 𝓓_*R*_1__, we have exhausted all substrings in 𝓓_*R*_2__ that are extendable to the left and whose extended part has edit distance lesser than or equal to 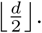 Therefore, we put a constraint on left extensions such that the edit distance of the extended part should be greater than 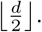

### Recursion

We recursively divide the read until the edit distance required for the alignment becomes zero. This corresponds to creating a binary tree such that in each step the given read and its distance are divided by two, c.f. Figure 3. Once we are at a leaf, an exact matching algorithm is called to obtain a list of candidate locations and then it will be used to construct the candidate locations of the parent node. At the end, all candidate locations will be obtained from merging lists from all leaves to the root node. It is worth mentioning that there exist numerous exact match finders [26, 27] that can be incorporated in the algorithm.

**Figure 3.**
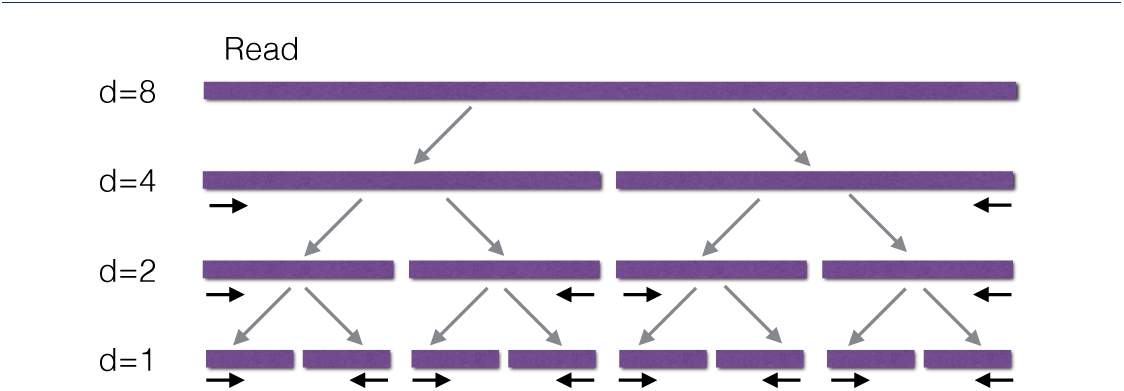
In this example a read with a tolerable edit distance *d* = 8 is given. The read is divided into two segments at the root. The left (respectively, right) segment is mapped to the reference genome with tolerable edit distance *d* = 4. Once a list of all genomic locations is obtained for the left (respectively, right) segment, extension to the right (respectively, left) is performed to find all the mappable locations on the genome with tolerable edit distance 8. The left and right segments are mapped through calling the procedure recursively.

### Extension: dynamic programming

Let *X*[*i*, *j*] denote a substring of *X* starting from *i*th location and ending at *j*th location. The basic dynamic programming to obtain the edit distance between two strings *U*[1: *n*] and *V*[1: *m*] creates a table of size *nm* where each cell stores a number *v*(*i*,*j*) representing the edit distance between *U*[1: *i*] and *V*[1: *j*].

In our algorithm, at each step a substring of read is needed to be extended and doubled to the left or right. The dynamic programming is started by feeding the first character to the left or right of the genomic substring and the first row is created. At each step, a new character is added and a new row is created. Since we have a constraint on the acceptable edit distance for the extension part, we terminate the procedure at any row if none of the values of the row satisfies the constraint. If the read can be extended, then the new genomic substring is added to the list of acceptable locations on the parent node.

We note that if there exists several location on the genome where a given substring of read *R* can be aligned to, then we can extend the substring over all such locations collectively. This fact can be used to speed up the algorithm. In particular, we create a tree where the root node corresponds to the initial row of the dynamic programming table. If the locations on the genome are extendable by any of the four DNA characters, then a child node is created for that character and a new row is added to the table, see Figure 4. The tree is extended until either the substring is doubled or the edit distance exceeds the given threshold *d*.

**Figure 4.**
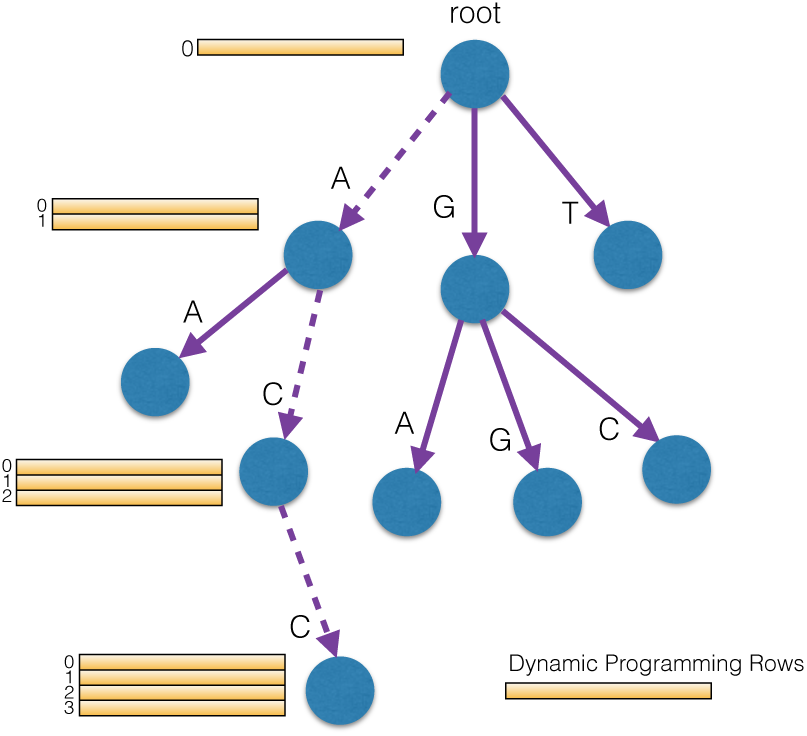
Here, the locations on the genome are extendable by three characters A,G, and T at the root. A new row is added to the dynamic programming table at each child of the root. The same procedure is performed at each internal node. In this way, the parent table is only constructed once throughout the algorithm.

### Heuristic methods for very long read alignment

In alignment of very long reads, it is not efficient and not required to align all the segment of the read to make sure that all possible candidate locations on the reference genome are identified. In fact, a few segments anchored reliably to the genome is enough for aligning the long reads to its true location. This idea is first presented in [14] where it is shown that a substring of length about 60-80 bases is statistically sufficient to anchor a long read to the reference genome. Incorporating similar ideas, we extend SureMap to handle long and very long reads.

Borrowing ideas from [14], a set of short substrings (seeds) are extracted from a read and aligned to the reference genome based on the methods presented in preceding sections. As early as three seeds can be mapped to the reference genome in such a way that the relative positions of the seeds on the read is almost the same as their mapped locations on the genome, then the whole read is anchored to the corresponding location.

After anchoring, it is desired to find the best global alignment between the read and the anchored location. However, the optimal procedure based on dynamic programming is tedious and time consuming. To overcome this, SureMap uses a simple heuristic algorithm to find a global alignment between two long strings which performs very well in practice.

The simple heuristic global alignment operates based on the following ideas. A read with length *L* is partitioned into 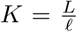 fragments of length *ℓ*. These fragments are denoted by *F*_1_, …, *F_K_*. Suppose *F*_1_, …, *F_k_* are aligned to the genome and the end position of the alignment is obtained. We call this position *e_k_*. Then, *F*_*k*+1_ can be aligned to the genome by assuming the starting position of the alignment is *e_k_*. In this way, we can obtain *e*_*k*+1_ and proceed to the next fragment. As it is observed experimentally, the end (starting) positions {*e*_1_, …, *e_K_*} may not be correct but the overall alignment is almost complete. To correct the end/starting positions of the read, say *e_k_*, we construct a fragment of length *ℓ* by concatenating the last 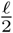 substring of *F_k_* with the first 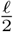 substring of *F*_*k*+1_. The obtained substring is re-aligned to the genome and possible corrections are made on the overall alignment.

## Data and software availability

The SureMap source code is freely available at: https://github.com/MohammadJRS/SureMap.

The software is provided under the MIT license, available at: https://github.com/MohammadJRS/SureMap/blob/master/LICENSE.

## Declarations

### Ethics approval and consent to participate

Not applicable

### Consent for publication

Not applicable

### Availability of data and materials section

- The datasets analysed during the current study are available in the NCBI SRA repository, SRR099983, ERR1676721
- The datasets analysed during the current study are available in the pacbio dataset repository, m130929-subread 1

### Authors’ contributions

MJ and ABL designed the general method for mapping, ABL choosed the samples for testing the software. MJ implemented the algorithms and performed the experiements. All authors wrote and approved the final manuscript.

